# The Time-Scale of Recombination Rate Evolution in Great Apes

**DOI:** 10.1101/013755

**Authors:** Laurie S Stevison, August E Woerner, Jeffrey M Kidd, Joanna L Kelley, Krishna R Veeramah, Kimberly F. McManus, Great Ape Genome Project, Carlos D Bustamante, Michael F Hammer, Jeffrey D Wall

## Abstract

We present three linkage-disequilibrium (LD)-based recombination maps generated using whole-genome sequencing data of 10 Nigerian chimpanzees, 13 bonobos, and 15 western gorillas, collected as part of the Great Ape Genome Project (Prado-Martinez et al. 2013). Using species-specific PRDM9 sequences to predict potential binding sites, we identified an important role for PRDM9 in predicting recombination rate variation broadly across great apes. Our results are contrary to previous research that PRDM9 is not associated with recombination in western chimpanzees (Auton et al. 2012). Additionally, we show that fewer hotspots are shared among chimpanzee subspecies than within human populations, further narrowing the time-scale of complete hotspot turnover. We quantified the variation in the biased distribution of recombination rates towards recombination hotspots across great apes. We found that correlations between broad-scale recombination rates decline more rapidly than nucleotide divergence between species. We also compared the skew of recombination rates at centromeres and telomeres between species and show a skew from chromosome means extending as far as 10-15 Mb from chromosome ends. Further, we examined broad-scale recombination rate changes near a translocation in gorillas and found minimal differences as compared to other great ape species perhaps because the coordinates relative to the chromosome ends were unaffected. Finally, based on multiple linear regression analysis, we found that various correlates of recombination rate persist throughout primates including repeats, diversity, divergence and local effective population size (N_e_). Our study is the first to analyze within-and between-species genome-wide recombination rate variation in several close relatives.

## Introduction

The time-scale over which recombination rates change is likely to vary from taxon to taxon. Several studies have noted the importance of scale when comparing results between studies and when comparing recombination rates within and between species (Stevison and Noor 2010; Auton et al. 2012; Chan et al. 2012). Comparison of recombination rates between close relatives has shown that recombination rates have rapid turnover on the scale of recombination hotspots (1-2 kb), but are correlated between species when examined at intervals of ~1 Mb (Serre et al. 2005; Duret and Arndt 2008; Laayouni et al. 2011). However, most previous between species comparisons have been focused on either very closely related taxa or distant relatives (Smukowski and Noor 2011). Nonetheless, differences in the conservation of recombination rates at various scales suggest different mechanisms control broad and fine-scale patterns of recombination rates across the genome. While recombination rates are free to evolve in different directions as species diverge, meiotic recombination is a tightly regulated cellular process and thus broad-scale rates may be limited both mechanistically and evolutionarily in how much they can change (Brooks 1988; Kauppi et al. 2004). Mechanistically, recombination is necessary to stabilize chromosomes during meiosis, but excessive recombination or errors in this pathway can lead to birth defects, disease, and/or various cancers (e.g. aneuploidy, Bloom syndrome or some breast/ovarian cancers). Evolutionarily, recombination helps to shuffle beneficial alleles onto common genetic backgrounds, facilitating the efficacy of selection. However, too much recombination can break down these associations.

Despite an often strong correlation of recombination rates between species at broad-scales, variation in the mechanisms controlling the distribution of recombination hotspots likely leads to the rapid turnover of fine-scale recombination rates. In bacteria hotspot determination is localized to χ sites, whereas in mammals, such as humans and mice, it has been shown that the transcription factor PRDM9 co-localizes with hotspots (Mihola et al. 2009; Baudat et al. 2010). However, the universal role of this protein remains unclear. For example, in dogs, the PRDM9 protein sequence is truncated and recombination hotspots are localized based on functional elements in the genome (Auton et al. 2013). In chimpanzees, there has thus far been no evidence that recombination rates are higher in regions with suspected PRDM9 binding (Auton et al. 2012). Unlike what has been shown in dogs, the chimpanzee PRDM9 protein is fully functional, and a recent survey of PRDM9 diversity in primates has shown pervasive diversifying selection for this protein throughout primates (Schwartz et al. 2014).

Contrary to strong positive selection of this protein in primates, comparisons of predicted binding motifs for the DNA binding zinc finger domain of PRDM9 have revealed short recurring submotifs among alleles from close relatives shared within populations of various primates (Myers et al. 2005; Auton et al. 2012; Schwartz et al. 2014). For example, in humans, two major submotifs of PRDM9 are associated with binding to recombination hotspots. This includes a 13-bp submotif CCnCCnTnnCCnC common to PRDM9 alleles found in European populations, and a 17-bp submotif CCnCnnTnnnCnTnnC more commonly associated with PRDM9 alleles found in African populations (Hinch et al. 2011a) (see Figure S8). For chimps and bonobos, four submotif regions, including a recently described internal submotif AnTTnnAnTCnTCC have been shown to recur in several alleles of PRDM9 across the *Pan* genus (Auton et al. 2012; Schwartz et al. 2014). The three submotifs searched in the recent chimpanzee recombination analysis include two motifs not present in western chimpanzees. One motif is predicted to be ancestral between eastern and western chimpanzee subspecies (E1) and the other is predicted to be ancestral between chimpanzees and bonobos (A1). For gorillas, an internal submotif CCnAnnCCTC was recently identified based on a smaller subset of PRDM9 alleles (Schwartz et al. 2014). The characterization of unique submotifs of PRDM9 to specific populations or subspecies further supports the rapid turnover of hotspots as species diverge.

Despite recent efforts to comprehensively sequence PRDM9 across primates, it remains to be seen whether the lack of PRDM9 localization to recombination hotspots in western chimpanzees is an exception among primates without corresponding recombination rate estimates. The first genome-wide recombination map in humans was a pedigree-based map of a European population (Broman et al. 1998). Since then, other pedigree-based maps of human populations have been generated for Europeans (Kong et al. 2002; Coop et al. 2008) and Asians (Bleazard et al. 2013). Genome-wide LD-based recombination maps have also been generated using HapMap genotype data, and include European (CEU), Yoruba (YRI), and Asian (CHB+JPT) population rate estimates (Myers et al. 2005). Recently, the 1000-genomes project constructed LD-based recombination maps from low-coverage whole-genome sequence data for the same populations (Altshuler et al. 2010). Another approach to fine-mapping recombination events in the genome has used local ancestry methods to build recombination maps for African-Americans (Hinch et al. 2011b; Wegmann et al. 2011). Finally, in 2012, the first non-human primate fine-scale recombination map was published using ten unrelated whole genome sequences of western chimpanzees (Auton et al. 2012). While there are now a number of population-specific recombination maps in humans and a western chimpanzee map (PanMap), recombination rate estimates outside of these groups are imperative to assess PRDM9’s role in hotspot determination. To gain a broader understanding of PRDM9’s role in recombination rates among great apes, we present three new LD-based recombination maps for Nigerian chimpanzees, bonobos and western gorillas collected as part of the Great Ape Genome Project (Prado-Martinez et al. 2013).

Because this dataset also represents a diverse group of closely related species with varying amounts of nucleotide divergence, we used it to address additional fundamental questions regarding the time-scale of recombination rate evolution. First, we sought to compare the distribution of recombination rate across the genomes of great apes. Previously, it has been shown that nearly 80% of recombination events occur in <20% of the physical sequence of the genome, localizing mostly to recombination hotspots (McVean et al. 2004). Further, recombination is more strongly biased towards hotspots in European recombination maps, but less so in African or chimpanzee recombination maps. For this analysis, we adopted the use of the Gini coefficient, which has been used in economics to compare the distribution of wealth among countries, and was recently applied to analysis of the cumulative distribution of recombination in *C. elegans* (Dorfman 1979; Kaur and Rockman 2014). In this context, it was shown to be a more robust way to directly compare the area under to curve of these cumulative distribution functions (Kaur and Rockman 2014). Secondly, we sought to compare broad-scale patterns of recombination rate divergence. This analysis included a comparison at various scales of the rate at which recombination rates diverge and nucleotide sequences divergence between species to understand the relative constraints on each. We also examined how large-scale chromosomal differences impact recombination rates. One major large-scale chromosomal change is the fusion of chromosomes 2a and 2b in the human lineage forming a much larger metacentric chromosome in humans. The previous fine-scale map of recombination in chimpanzee revealed the centromere of the now human chromosome 2 has much higher recombination rates in chimpanzees, where it is the telomeres of chromosomes 2a and 2b (Auton et al. 2012). We also examined the skew in recombination rates typically present in telomeric and centromeric regions.

Finally, because recombination rates have been shown to correlate with genetic features such as polymorphism, divergence, GC-content, repeat content and specific sequence motifs (Jensen-Seaman et al. 2004; Coop and Przeworski 2007; Stevison and Noor 2010), we sought to determine the amount of variation in recombination rate that can be explained by various genetic features. The finding that recombination rate variation across the genome correlates with a variety of genetic features has led many to attempt to determine which evolutionary forces drive these associations. For example, many studies have found that recombination-mediated linked selection drives the ubiquitous correlation observed between recombination rate and nucleotide polymorphism in many taxa (Begun and Aquadro 1992; McGaugh et al. 2012; Webster and Hurst 2012). Alternatively, a biased mutation process known as GC-biased gene conversion (gBGC) has been shown to explain the correlation between GC-content and recombination in most cases (Marais et al. 2001; Birdsell 2002; Marais 2003; Duret and Galtier 2009; Galtier et al. 2009), but see (Hey and Kliman 2002; Kliman and Hey 2003). For this analysis, we comprehensively represented the various genetic feature data available in primates and sought to normalize the explained variation by employing a multiple linear regression framework. In addition to the findings we present here, we anticipate this resource will be useful both for imputing and phasing future genotype data collected and in uncovering unique patterns of selection and demography in these species (McManus et al. *In press*).

## Results

We used 10 *Pan troglodytes ellioti*, 13 *Pan paniscus*, and 15 *Gorilla gorilla gorilla* (**Table S1**) whole-genomes sequences (mean coverage 26.34x) to construct recombination maps for each species, using a similar approach to the western chimpanzee map (Auton et al. 2012). The final maps were constructed from 4.2, 8.5, and 7.8 million SNPs for bonobo, Nigerian chimpanzee, and western gorilla, respectively, as compared to 5.3 and 1.6 million sites used in the western chimpanzee and HapMap projects, respectively. The genome-wide average recombination rates were 0.641, 0.8, and 0.944 ρ/kb in bonobo, Nigerian chimpanzee, and western gorilla respectively. For a robust comparison between our maps and existing human and western chimpanzee maps, we identified blocks for each non-human genome that were syntenic with human (**Figure S2-4**) binned to 1 Mb (**Table S2**; Figure 1), 500 kb and 100 kb. Additionally, Figure S5 shows a plot of recombination rates across the genomes of the great apes compared here. To identify recombination hotspot locations, we implemented a version of LDhot that follows the approach of Myers *et al*. (2005). Briefly, for a 20 kb region, we used a likelihood ratio test to determine whether the central 2 kb had an elevated recombination rate relative to the surrounding sequence (see **Figure S6** for comparison to other methods). Using this approach, we identified 10704, 8037, and 22012 hotspots in bonobo, Nigerian chimpanzee, and western gorilla, respectively.

**Fig. 1.**
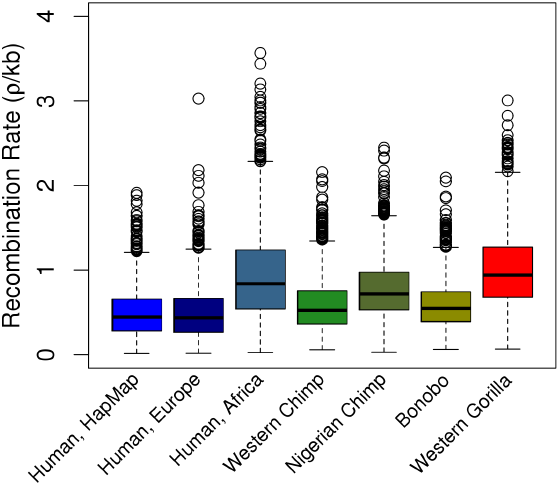
Broad-scale comparisons of recombination rates across great apes. A comparison of the mean and variance of average recombination rates in 1Mb bins of previously published HapMap, European and African human populations from Auton *et al*. (2012) combined with the Nigerian chimpanzee, bonobo and western gorilla recombination rates generated in this study.

### Localization of PRDM9 to hotspot regions

Together with the publicly available hotspots from HapMap (Myers et al. 2005) and western chimpanzee (Auton et al. 2012), we investigated the importance of various PRDM9 submotifs in localizing to population-specific hotspots by calculating position weight matrices (PWMs) identified based on species-specific protein sequences of PRDM9 (**Figure S8**). We then compared the proportion of hotspots versus matched coldspots within 100 kb with a PBS (**Table 1**) similar to an approach used in a recent mouse study (Brunschwig et al. 2012). We also summed the total potential binding sites (PBS) count across all hotspots and coldspots over all submotifs and found significant enrichment of both proportion of hotspots with a PBS and total motif count in the hotspots as compared to coldspots for all species, except when using our newly generated hotspots for western chimpanzee. Both human submotifs were significantly associated with higher PBS counts and proportions in hotspots versus coldspots, though allele C was marginally significant in CEU hotspots. Using the western chimpanzee hotspots from Auton *et al*. 2012, we found that the Schwartz submotif, the western chimpanzee-specific submotif, and to a lesser extent the A1 ancestral submotif were significantly associated with more PBS counts in hotspots versus coldspots. This signal mostly went away when using our newly generated set of hotspots in the western chimpanzee, though the Schwartz submotif remained significantly associated. The Nigerian chimpanzee showed significance for the western-specific submotif, the E1 ancestral submotif, and the Schwartz submotif. The lack of association with A1 in Nigerian suggests this allele may not be segregating in this population, but direct sequencing of PRDM9 alleles in this population would address this question more directly. Bonobo showed a weak association with the A1, E1 and Schwartz submotifs, but not the western chimpanzee specific submotif, which is not surprising as this is most likely a derived allele of PRDM9. Finally, gorillas showed a strong association with both the Schwartz submotif and the additional ‘motif 2’ submotif we analyzed here.

We further analyzed for each submotif of PRDM9, the PBS count relative to hotspot strength (**Figure S9**) and percentage GC (**Figure S10**) in hotspots versus coldspot regions. Based on partitioning of hotspots based on their intensities, we found that for moderate and strong human hotspots, the number of binding sites is significantly higher than matched coldspots in humans for human allele A in both CEU and YRI and for human allele C in YRI only (**Figure S9A**). For western chimpanzee, the increased association with stronger hotspots is only apparent when using the newly generated set of hotspots (**Figure S9C**) for the submotifs significant in Table 1. The Nigerian chimpanzee results mostly matched those in Tables 1 and S3; however, the submotif A1 shows a larger difference between hotspots and coldspots in the highest two quartiles for hotspot strength, suggesting some historical signature remains at least for the strongest hotspots. We found the putatively ancestral PRDM9 submotif, A1, not significant overall in chimpanzee, to have higher hotspot motif counts in stronger hotspots. The Bonobo and Gorilla results for this analysis matched the results in Table 1. When we broke the results up based on GC content, the human alleles have high GC content and the majority of binding sites correspond the highest GC bin. For allele A, the pattern of a higher number of PBS in hotspots versus coldspots persists across all bins; however, for allele C, the PBS count is similar between the two regions for the highest bin, suggesting that GC count is not driving this result for either submotif. For western chimpanzee, the Schwartz submotif does not show a difference in the lowest GC bins for the PanMap hotspots, but this difference is apparent with the newly generated set of hotspots. Further, the newly generated set of hotspots revealed that for the western-specific and A1 submotifs, the lower GC bins show the strongest pattern, further suggesting that GC content is not driving this result. For Bonobo, the E1 submotif does have a stronger signal in higher GC bins, suggesting this result is less likely to be truly associated with higher recombination. For gorilla, there appears to be a consistent difference across all bins for both submotifs.

**Table 1.** PRDM9 Summary. Comparison between identified hotspots and matched cold spot regions based on proximity, size and GC-content. For each set of hotspots, the proportion of hotspots versus cold spots that contain a predicted binding sequence (PBS) based on species-specific position weight matrices (PWM) of the PRDM9 sequence. This is contrasted on the right with the total predicted binding motifs present in the combined sets of hotspots versus cold spots. Results for each relevant submotif and the combined results are presented for each genetic map with the source of the hotspots listed.

| Recombination Data | Submotif | Proportion of regions with a PBS |  | Binomial test p-value | Total motif count |  | Binomial test p-value | Hotspot source |
| --- | --- | --- | --- | --- | --- | --- | --- | --- |
|  |  | Hotspots | Cold spots |  | Hotspots | Cold spots |  |  |
| HapMap CEU |  | 0.50 | 0.45 | **2.70E-16** | 41984 | 39168 | **4.95E-23** | HapMap |
|  | Allele A | 0.38 | 0.33 | **2.23E-23** | 27808 | 25390 | **1.06E-25** |  |
|  | Allele C | 0.29 | 0.27 | **1.29E-06** | 14176 | 13778 | **0.02** |  |
| HapMap YRI |  | 0.49 | 0.44 | **7.71E-22** | 40732 | 37687 | **1.58E-27** | HapMap |
|  | Allele A | 0.38 | 0.32 | **9.04E-29** | 26901 | 24346 | **1.57E-29** |  |
|  | Allele C | 0.28 | 0.26 | **7.77E-07** | 13831 | 13341 | **3.01E-03** |  |
| Western chimpanzee |  | 0.64 | 0.60 | **0.02** | 5776 | 5480 | **0.01** | PanMap |
|  | Western | 0.28 | 0.27 | 0.26 | 1615 | 1502 | **0.04** |  |
|  | A1 | 0.21 | 0.19 | **0.04** | 1143 | 1042 | **0.03** |  |
|  | E1 | 0.26 | 0.26 | 0.93 | 1427 | 1499 | 0.19 |  |
|  | Schwartz | 0.28 | 0.25 | **8.04E-04** | 1591 | 1437 | **0.01** |  |
| Western chimpanzee |  | 0.60 | 0.59 | 0.62 | 12599 | 12410 | 0.23 | This study |
|  | Western | 0.27 | 0.28 | 0.26 | 3393 | 3534 | 0.09 |  |
|  | A1 | 0.21 | 0.21 | 0.92 | 2490 | 2496 | 0.94 |  |
|  | E1 | 0.25 | 0.26 | 0.13 | 3163 | 3241 | 0.34 |  |
|  | Schwartz | 0.28 | 0.25 | **1.61E-03** | 3553 | 3139 | **4.41E-07** |  |
| Nigerian chimpanzee |  | 0.63 | 0.58 | **2.55E-04** | 9316 | 8520 | **2.62E-09** | This study |
|  | Western | 0.29 | 0.26 | **3.85E-05** | 2654 | 2356 | **2.70E-05** |  |
|  | A1 | 0.21 | 0.20 | 0.29 | 1807 | 1745 | 0.31 |  |
|  | E1 | 0.27 | 0.25 | **0.01** | 2413 | 2214 | **3.60E-03** |  |
|  | Schwartz | 0.27 | 0.25 | **0.01** | 2442 | 2205 | **5.35E-04** |  |
| Bonobo |  | 0.61 | 0.59 | **0.05** | 14081 | 13369 | **1.77E-05** | This study |
|  | Western | 0.29 | 0.28 | 0.08 | 3968 | 3859 | 0.22 |  |
|  | A1 | 0.23 | 0.21 | **0.01** | 2928 | 2739 | **0.01** |  |
|  | E1 | 0.27 | 0.25 | **0.04** | 3556 | 3357 | **0.02** |  |
|  | Schwartz | 0.27 | 0.26 | **0.04** | 3629 | 3414 | **0.01** |  |
| Gorilla |  | 0.32 | 0.30 | **1.39E-05** | 10384 | 9421 | **8.08E-12** | This study |
|  | Schwartz | 0.23 | 0.21 | **1.94E-03** | 6429 | 6080 | **1.86E-03** |  |
|  | This study | 0.15 | 0.13 | **2.49E-06** | 3955 | 3341 | **6.93E-13** |  |

For a more direct comparison to the results of Auton *et al*. (2012), we also performed a genome scan for each submotif and summed the results over all submotifs for each group and compared the results with a corresponding null motif (**Figure S11**). For human, we show higher rates compared to the null for allele A in both CEU and YRI and for allele C in YRI only. Though the null rate for YRI in allele C is also locally high, this is perhaps due to our null motif being a random shuffling of the same PWM matrix. The genome-wide scan did not reveal any significant association between recombination rate and the submotifs in Nigerian chimpanzee and bonobo, similar to previous results in western chimpanzee. For gorilla, we found a higher recombination rate at motif 2, but not the Schwartz motif. We further split the genome-wide data based on whether each 1kb region had 0, 1 or 2+ predicted binding sites and further split these into GC quantiles as was done for the hotspot/coldspot analysis (Figure S12). For simplicity, we summed these results across all submotifs. We found that the recombination rate of both human and gorilla is higher in regions with higher PBS counts, and that this difference is consistent across GC bins and stronger for the true motifs versus the null motifs. These results were not observed in the *Pan* species, consistent with the full genome-wide search results.

In addition to examining PRDM9’s association with recombination rate, we examined the degree of overlap for recombination rate at hotspots across great apes (Figure 2). We found more overlap between human populations for recombination rates at hotspots identified in other populations than within western and Nigerian chimpanzee subspecies. It is worth noting that Figure 2A over-represents the amount of hotspots shared between human populations because the HapMap hotspots are a composite of population-specific hotspots from European, Yoruba and Asian populations. Therefore, we compared our population-specific hotspots results for chimps (Figure 2B & C) to the population-specific hotspot plots of European and Yoruba human populations in Auton et al (2012) supplementary Figure 6. Still, the degree of overlap in hotspots is much higher than Figure 2B & C. Another potential contribution to differences in the results is our method for identifying hotspots was slightly different than the methods used for both HapMap and western chimpanzee. However, we estimated hotspots in western chimpanzee using our method and did not see any qualitative difference in the degree of sharing with other populations (**Figure S7**). We found a similar amount of sharing between Nigerian chimpanzee and bonobo, though no sharing between bonobo and western chimpanzee (Figure 2D). We found no overlap of hotspots between gorilla and the other species studied here (Figure 2E), which is most likely due to a lack of any close relatives for comparison and the rapid turnover of hotspots.

**Fig. 2.**
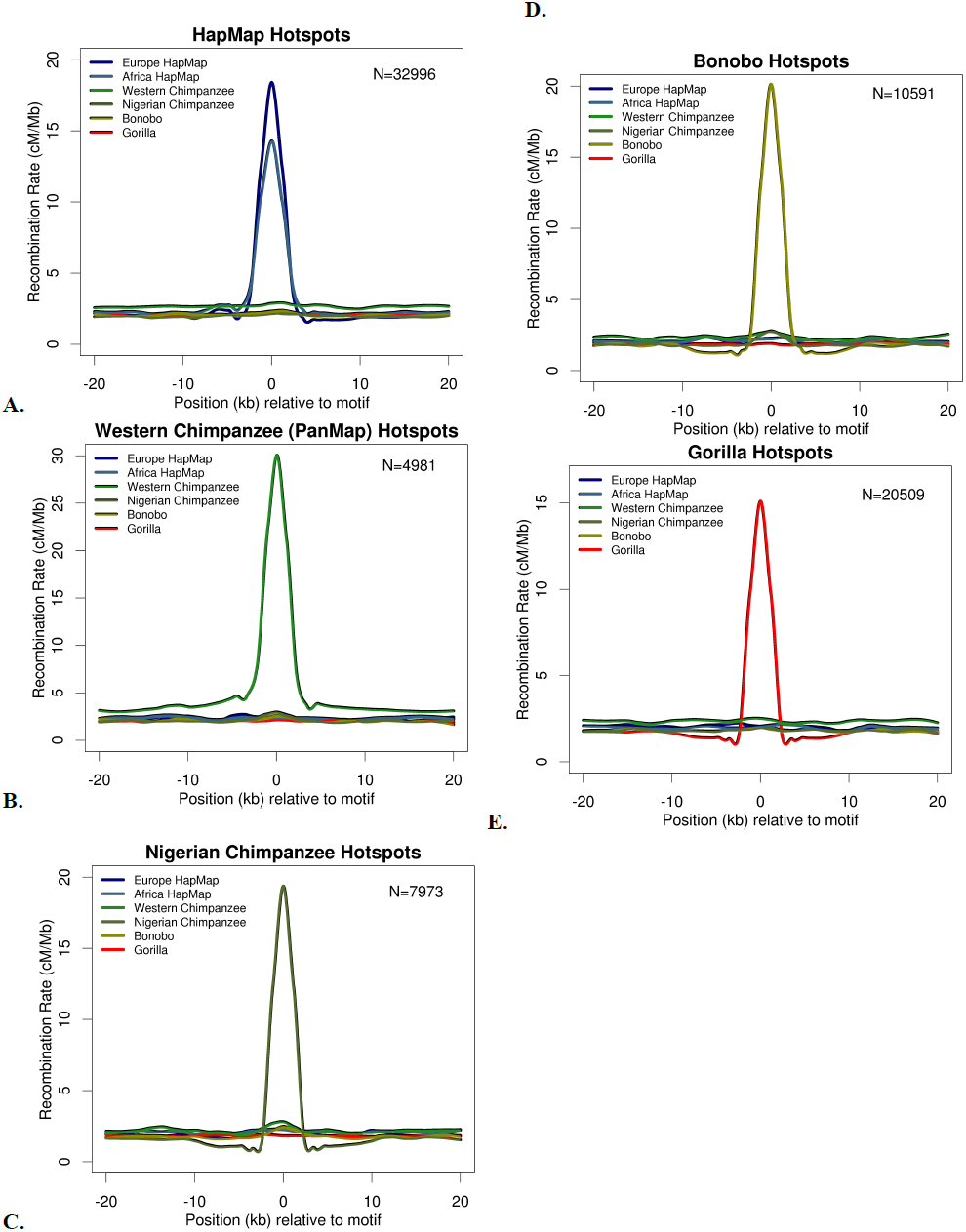
Recombination enrichment at species-specific hotspots. Degree of hotspot sharing and recombination rates for all maps in species-specific hotspots. Recombination rates for all maps 20kb upstream and downstream of (**A**) human hotspots from HapMap, (**B**) western chimpanzee hotspot centers from PanMap, (**C**) Nigerian chimpanzee hotspot centers, (**D**) bonobo hotspot centers and (**E**) western gorilla hotspot centers. Rates shown in cM/Mb and numbers of hotspots correspond to the number that mapped to hg18 genome.

### Distribution of recombination rate across genome

Another way to compare the recombination rates within and between great apes is to look at the distribution of recombination rates across the genome. Similar to previous studies, we found a biased recombination rate distribution whereby the majority of recombination (~75%) occurs in a fraction of the physical genome (~20%). From the corresponding Gini coefficient using the AUC of the Lorenz curve (Figure 3A), we show the European human population presents the strongest hotspot usage across the genome, similar to previous studies (Figure 3B). We confirm the differences between the CEU (Gini=0.771), YRI (Gini=0.688) and Chimp (Gini=0.677) maps previously published. For the new maps, we calculate a Gini coefficient that lies within the values of the extremes of these previously published maps. Specifically, we calculate a Gini coefficient of 0.704 for Gorilla, 0.713 for Bonobo and 0.726 for Nigerian Chimp. Using this statistic, we are able to show that the distribution of recombination rate across the genome is variable in great apes. The values from the recently published survey of Gini coefficients across various species were included as a reference point (Kaur and Rockman 2014)

**Fig. 3.**
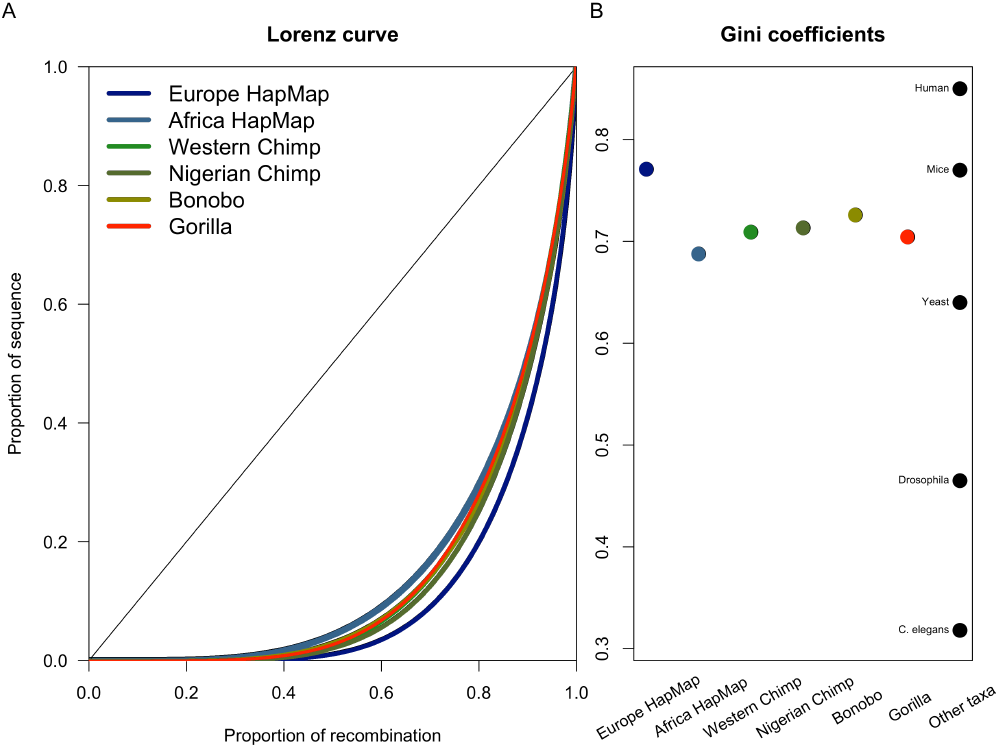
Genome-wide distribution of recombination rates. Cumulative distribution, or Lorenz curve of recombination rate plotted as proportion of recombination versus sequence for each recombination map (**A**). The diagonal represents a uniform distribution. Gini coefficients for each population map, and for comparison, other taxa reported in (Kaur and Rockman 2014), including a human estimate from (Kong et al. 2002) (**B**).

### Broad-scale comparisons

We calculated the Spearman rank correlation coefficient between all pairwise recombination rates and compared the correlation coefficients to the amount of nucleotide sequence divergence between each pair at 1 Mb (Figure 4), 500 kb (**Figure S14A**) and 100 kb (**Figure S14B**). We found that pairs with low nucleotide divergence experience a rapid decline in recombination rate correlation, whereas comparisons with higher nucleotide divergence have relatively similar recombination rate correlations. Further, the correlation at decreasing scales suggests that sequence divergence explains less of the variance in recombination rates at finer scales as has been previously shown.

**Figure 4.**
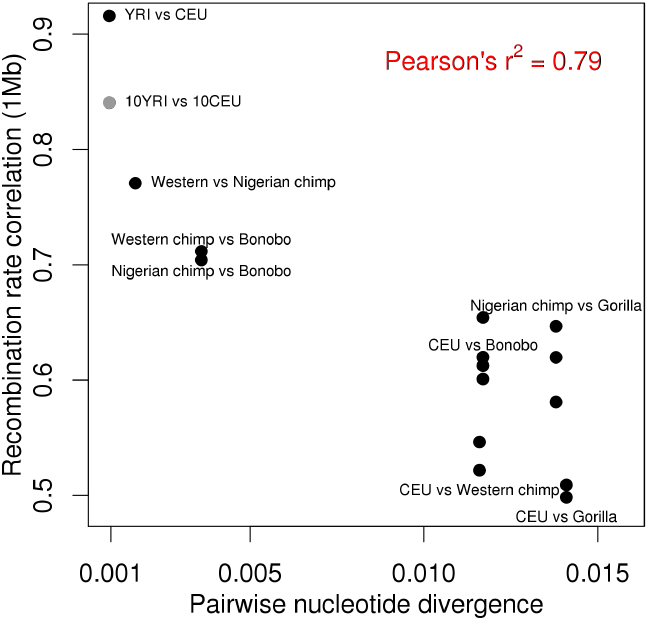
Recombination rate versus nucleotide divergence. Spearman rank correlation coefficient between all recombination maps at 1Mb (y-axis) versus the pairwise nucleotide divergence between pairs (x-axis) with various comparisons labeled.

Another interesting broad-scale recombination pattern is the skew in recombination rates at the ends and near the center of chromosomes. We quantified the extent of this skew across species, controlling for differences in recombination rate in each chromosome (**Figure S15**). While centromeric regions recovered to the mean of the chromosome within 5 Mb of the centromere, the skew at telomeres was more pronounced and continued for nearly 15 Mb from the chromosome end. We further looked at large-scale chromosomal changes across great apes, including the chromosome 2 fusion in humans and the chromosome 5/17 translocation in gorillas. We found that other non-human primates also have high recombination rates across the junction of chromosomes 2a/2b supporting its historical telomeric origin. However, bonobos have lower recombination rates across this region similar to what has been seen in humans (**Figure S16A**). Though this is most likely due to reduced coverage in this area for bonobos. We further found that the translocation event in gorillas did not influence broad-scale recombination rates, likely because it did not involve centromeric or telomeric regions (**Figure S16B & C**).

### Multiple linear regression analysis

We used a multiple linear regression analysis to evaluate the relationship between recombination and various genetic features. Despite controlling for various classes of repeats and selective elements, there is a strong positive correlation between both divergence, as measured by the distance to the ancestor of humans and orangutans and local N_e_ (as measured by pairwise nucleotide variation within species, π divided by pairwise divergence between species, D) with local recombination rate (Figure 5 and Figure S13). We show that local N_e_ as a predictor of recombination rate variation is most pronounced in species with large effective population size. We also show that GC content correlates with local recombination rate and is more pronounced in comparisons that are closely related to humans, perhaps owing to use of human GC content (Munch et al. 2014).

**Figure 5.**
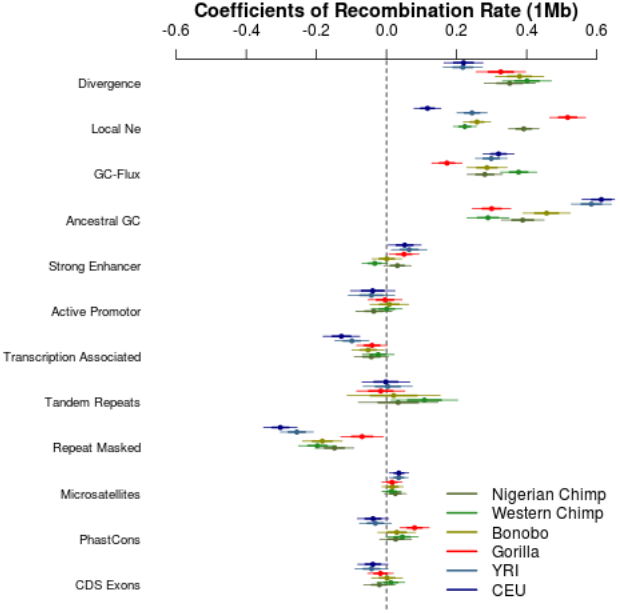
Genetic correlates of recombination rate. Multiple linear regression analysis results with normalized coefficients to compare relative weights of each genetic feature in contributing to the variance of recombination rate at 1 Mb.

## Discussion

By extending the number of whole-genome fine-scale genetic maps, we present a major advance in understanding recombination rate evolution across great apes. First, we have shown that PRDM9 explains recombination rate distribution across great apes, not just in humans as has been shown previously.

Crucial to our finding was the focused analysis at recombination hotspots rather than a wholegenome approach as has previously been used in this context. This analysis requires highquality recombination estimates at fine-scales. While we report significant enrichment of putative PRDM9 binding in recombination hotspot regions as compared with coldspot regions, we did not observe a significant association between PRDM9 binding broadly across the genome and local increases in recombination rates. Below we discuss some of the reasons for the incongruence between the genome-wide results and the hotspot results. Additionally, we have narrowed the time period of complete hotspot turnover between species to the split times between chimpanzees and bonobos. Further, we have analyzed broad scale patterns of recombination rate evolution and shown that the divergence in recombination rates between species occurs very rapidly when compared to nucleotide sequence divergence, and perhaps more closely tracks population divergence times. We show that large-scale chromosomal differences not involving chromosome ends are less likely to impact broad-scale recombination rates. Finally, we show that a subset of genetic features explains the majority of the variation in recombination rate and that the amount of variation explained depends in part on the effective population size.

### PRDM9 ubiquitously associates with great ape hotspots

As shown in Table 1, we find strong evidence that PRDM9 likely binds to recombination hotspot regions predominantly as compared to coldspot regions broadly across great apes. In fact for all other groups of great apes, we find at least one submotif of PRDM9 enriched across hotspot regions as compared to coldspot regions. Because the signal in western chimpanzees is mainly reflected in the newly identified internal submotif, it is not surprising that the earlier study in this group failed to identify an important role for PRDM9 in recombination rate association. Indeed the western chimpanzee specific submotif of PRDM9 is most strongly associated with predicted binding in Nigerian chimpanzee recombination hotspots. This is possibly due to shared alleles of PRDM9 between these two subspecies. However, the Nigerian chimpanzees have not been previously included in surveys of PRDM9 diversity. Further, the A1 ancestral motif does not seem to be active in this group despite its activity in bonobos supporting the rapid turnover of hotspot landscapes observed between these groups. Additionally, the PRDM9 submotif which is putatively ancestral across chimpanzee subspecies seems to be active in the outgroup of bonobo, suggesting it appears in other older alleles of PRDM9 present in bonobos as well. By breaking hotspots into groups with potential binding of various submotifs of PRDM9, we can break down the hotspot landscape. This supports recent evidence in humans that LD-based recombination rate estimates represent a composite landscape with distinct landscapes superimposed to yield a population average (Pratto et al. 2014).

In the western chimpanzee study examining the association between PRDM9 binding and recombination rates, they searched for computationally predicted binding of PRDM9 submotifs across the whole genome. Using the top hits where PRDM9 was predicted to bind, they examined recombination rates as compared to a motif with a single site swapped for another. Because we were interested in colocalization of recombination and PRDM9 and we already know where recombination hotspots are in the genome, we chose instead to focus on hotspot regions. Additionally, computational methods to predict transcription factor binding in the genome can be unreliable (Billings et al. 2013), especially in the absence of specific information about accessible DNA during the time period when the transcription factor is most active (during meiosis in the case of PRDM9). This can lead to a lot of false positives for regions of the genome that may not be accessible to the protein when it is active. For example, there is no data on the state of open chromatin for the specific time period when recombination occurs for these taxa to give an idea of the accessibility of the DNA to the PRDM9 protein during the period when it is most active. Additionally, there may be putatively pleiotropic functions of PRDM9 that potentially involve DNA binding, but do not play a role in initiating recombination. Finally, the recent diversity survey of PRDM9 in chimpanzees also searched for the major submotifs across the human, chimpanzee and gorilla genomes and found they were equally prevalent across all genomes and highly abundant (Schwartz et al. 2014). Nonetheless, to compare our results to those of Auton *et al*. (2012), we further examined rate differences associated with PBSs along the genome irrespective of local recombination rate. We compared recombination rates in regions with a PBS based on the species-specific PRDM9 PWM versus a null version generated by shuffling the original PWM. We found higher recombination rates near PRDM9 PBSs versus the null PBS for gorilla, similar to humans (**Figure S11**), and irrespective of GC content (**Figure S12**), but not for *Pan* species. We attribute this to the loss of sensitivity of a genomewide search in the *Pan* group due to the high diversity at the PRDM9 locus in the *Pan* genus (Schwartz et al. 2014). The higher allelic diversity at PRDM9, especially in *Pan* species, likely contributes to population rate estimates based on patterns of LD being a composite of multiple distinct hotspot landscapes superimposed (Pratto et al. 2014). Further, LD-based maps are less likely to reflect recent changes in the recombination landscape and the rapid turnover of hotspots in this group may also render a genome-wide approach difficult in uncovering a true association between specific PRDM9 submotifs and recombination rates. In human and gorilla, both the age of the alleles and the frequency in the population sampled likely yield higher power to detect such a signal, which may also explain the reduced signal for both the Allele C and the gorilla Schwartz submotifs. However, a lack of a signal only reflects our inability to detect it and not necessarily its true absence in these groups.

We found that variation in the great ape recombination maps presented here are similar to other vertebrate taxa with functional PRDM9 recently surveyed (Schwartz et al. 2014). These results are consistent with previous work reporting a dominant PRDM9 allele for determining hotspot locations across the genome in European populations, driving the bias towards hotspot usage (Altshuler et al.

2010). Further, higher allelic diversity and levels of within population heterozygosity of PRDM9 likely contribute to a more even distribution of recombination rates across the genome, with distinct hotspot landscapes averaged over the longer population history of chimpanzee, bonobo and gorilla (Berg et al. 2011; Schwartz et al. 2014). However, because the amount of variation in Gini coefficients across great apes is rather small, it would be difficult to distinguish between the impact of PRDM9 variation and variation in effective population size (N_e_) (Auton et al. 2013).

### Within and between species comparison of recombination rate evolution

Access to multiple within-and between-species comparisons across great apes allowed us to perform a comprehensive comparison of recombination rate variation at fine-scales. The two chimpanzee subspecies diverged ~400-600 kya (Bowden et al. 2012), whereas the two human populations diverged ~50-60 kya, indicating a very rapid change in the hotspot landscape over a short evolutionary time period. Given recent reports of ~150 overlapping hotspots between chimpanzees and human on chromosome 21 (Wang and Rannala 2014), the lack of overlap in recombination rates at hotspots between the two chimpanzee subspecies is surprising (Figure 2B&C). The lower coverage of individuals (mean 9.45x coverage) and the high error rate (34-41%) at the fine-scale (<10 kb) reported in the western chimpanzee study indicates that individual SNP calls may not be accurate and direct comparisons with western chimpanzee may not be appropriate (Auton et al. 2012). Nonetheless, because we find a similar lack of shared hotspots between bonobo and chimpanzee subspecies (Figure 2D), complete hotspot turnover occurs within the boundaries of the chimpanzee subspecies divergence time (~0.5 mya) and the chimpanzee-bonobo divergence time (~1-2 mya).

### Broad-scale recombination rate changes occur more rapidly than nucleotide divergence

Because fine-scale recombination rates change rapidly within species, focusing on broad-scale recombination rate changes between species allowed us to identify changes that occurred over longer evolutionary time frames. These results suggest that the amount of change in recombination rate over time plateaus after a few millions years with gorilla versus human comparisons largely overlapping chimpanzee versus human (Figure 4 and Figure S14).

We further analyzed the skew of recombination rates at chromosome ends across great apes and found a stronger skew at telomeric regions that centromeric regions. Previous studies account for this skew by removing ~5-10 Mb nearby centromeres and telomeres (Serre et al. 2005), which while sufficient for centromeres, but may not adequately account for the telomeric skew in great apes. We also plotted recombination rates across the chromosome 5/17 translocation present in gorillas and found that unlike the chromosome 2 fusion event, this does not seem to disrupt the broad-scale recombination landscape (**Figure S16B & C**). This is perhaps due to the fact that this change does not involve any chromosome end regions where rates have broad-scale differences.

While we examined many possible factors which may be associated with recombination rates, we found that variance was mostly explained by divergences, local N_e_ and GC content. While the relationship with local N_e_ is not surprising, the observed association between divergence and recombination rate is unexpected. Although this pattern should indicate recombination to be a mutagenic process, recent results in humans strongly discount this possibility (Schaibley et al. 2013). An alternative explanation is that functional density (regions subject to selection) is non-randomly distributed and perhaps is higher in low recombining regions (Cutter and Payseur 2013). These results suggest that despite rapid turnover in local recombination rates correlations between specific genetic features and recombination rates are consistent across great apes, but the degree is dependent on factors such as N_e_ and population divergence.

## Methods

### Samples, Sequencing and SNP calling

Samples for the fine-scale recombination maps presented here were collected and described in the Great Ape Genome Diversity Consortium (Prado-Martinez et al. 2013). From the 88 samples described, 38 were used here to estimate genome-wide recombination rates for three populations of three major species: 13 bonobos without known geographical origin (*Pan paniscus*); 10 chimpanzees from Nigeria (*Pan troglodytes ellioti*); 15 western gorillas from Cameroon and Congo (*Gorilla gorilla gorilla*). This subset of individuals is described in **Table S1**.

A detailed description of sequencing, reference mapping and SNP/Variant calling can be found in Prado-Martinez *et al* 2013. Briefly, samples were sequenced on an Illumina sequencing platform (HiSeq 2000) with data production at four different sequencing centers, sequence reads were mapped to both the human reference genome (hg18) and each species-specific reference (PanTro 2.1.4, Ensembl release 65 for *Pan* and gorGor3, Ensembl release 62 for *Gorilla*). SNP calling was performed using the Genome Analysis Toolkit (GATK) software (DePristo et al. 2011). Final coverage for the individuals used here is included in **Table S1**.

### Data filtering and synteny designation

The process of data filtering and rate estimated was carefully matched to be similar to that used in the recent PanMap project (Auton et al. 2012). Using both the human-based mapping and the species-specific mapping of the reads for each species, the data were filtered using a combination of VCFtools (Danecek et al. 2011) and custom scripts (Stevison 2015). First, sites with more than 80% missing data were removed and sites with a minor allele count less than one were removed to avoid fixed differences (specifically to the human reference) or variants segregating in samples not included here. Then, sites within 15bp of each other were thinned to only retain a single site. Next, each site was reciprocally liftedover using the UCSC tool liftOver (minMatch=0.1) (Hinrichs et al. 2006). Filtering of the mapped sites following the reciprocal liftover process retained only sites that mapped back to the same position as the original input file. Then sites not in Hardy-Weinberg were removed (cutoff=0.001). After these series of filters were imposed on both sets of SNP calls, the remaining sites were intersected using the liftover positions between the two assemblies. From then on the species-specific orientation was used to maintain the inherited order of sites for subsequent computational phasing and recombination rate estimation steps.

Using the intersected set of sites, synteny blocks were defined based on the coordinates in both the human and non-human primate reference genomes. Then, using these matched sets of coordinates, consecutive pairs of sites were considered along the sequence of each chromosome. Both orientation, based on whether coordinates were increasing or decreasing relative to each genome, and a maximum distance of 50kb were used to define whether they will be included in the current block or start a new block. Distance calculations were performed using coordinates in both references genome so that gaps in either genome would terminate the block.

Once syntenic blocks were defined and sites between blocks were eliminated, adjacent blocks were considered for concatenation in order to reduce the overall number of blocks. This procedure involved concatenating blocks only if it involved removing no more than 300 intervening sites and the distance between all adjacent sites was maintained to be below 50kb. Additionally, syntenic blocks below 300 sites were removed in this step due to this being the minimum number of sites required to estimate rates using LDhat. Briefly, the LDhat software performs rate estimation considering 100 sites at a time.

Further, as a standard practice, the 100 bounding sites are typically discarded due to edge effects. Based on this information, it was determined that any blocks below 300 sites would not provide accurate rate estimates, and that ultimately only the central 100 sites would be used for reporting rate estimates. The process of concatenating blocks was repeated to further reduce the number of blocks, while again ensuring that blocks maintain the same orientation and gaps greater than 50kb were not observed between any pair of adjacent sites. See **Figure S1** for the distribution of block sizes for each species. See **Figures S2-4** for plots of each set of coordinates as mapped in the human reference versus the non-human reference genome, highlighting large-scale differences from human in orientation for each species.

### Phasing, imputation and recombination rate estimation

Computational phasing and imputation to infer bases at missing sites was performed on each syntenic block using the software fastPHASE v1.2 (Scheet and Stephens 2006) using the option K=10 for the number of haplotype clusters, which was the optimum after running the full cross validation on a subset of the data. While other comparisons have found the K=20 option to be more accurate, these were based on samples size of ∽100 (Browning and Browning 2011). Here, our sample size was much smaller, so we performed validation on the largest sample size using the gorilla data, N=15.

For improved phasing accuracy, the variants were re-phased using PHASE v2.1 (Stephens and Donnelly 2003) as described in (Auton et al. 2012). Briefly, the synteny blocks were split into 400 SNPs with 100 SNP overlap between files. PHASE was run on each file separately and synteny blocks were pieced back together using the minimum hamming distance based on the 100 overlapping SNPs. The rephased output was converted back to a vcf file. An additional filter based on minor allele frequency was performed afterwards (cutoff=0.05). If the resulting syntenic block still had a minimum of 300 sites, it was further split into 4000 SNP blocks with 100 SNP overlap and converted to input for the software LDhat v2.1 (Fearnhead and Donnelly 2001; International HapMap 2005), otherwise it was removed from further analysis. LDhat was run for 60 million iterations with a block penalty of 5, sampling every 40,000 steps (see Auton et al. 2012). After rate estimation, synteny blocks were pasted back into continuous segments while removing the 100 SNP edges between independent runs (Myers et al. 2005). All blocks in a single chromosome were also written to a single file, though the rate estimates are not continuous across the whole chromosome.

### Comparisons to existing maps

To get comparable recombination rate data for the published human and western chimp maps, the ‘syntenic genetic map’ file was downloaded from the *‘haplotypes*’ directory on ftp://birch.well.ox.ac.uk/panMap/. Rate estimates for western chimps (panTro2_map column), HapMap_CEU, and HapMap_YRI were extracted. Rates were converted to ρ for human maps using N_e_=10040 for CEU and N_e_=19064 for YRI (Auton et al. 2012), and all rates were scaled according to physical distance in kb. Additionally, HapMap rates in hg19 were downloaded from ftp://ftp.ncbi.nlm.nih.gov/hapmap/recombination/2011-01_phaseII_B37/. Further, our map estimates were converted from ρ/kb to cM/Mb following the approach of McVean et al (2004). We made the simplifying assumption that the average probability of recombination per bp is exactly the same across taxa. While the assumption of uniformity in map lengths is not ideal, the timescale associated with the N_e_ used in genetic maps is not known, and while the regression approach of (Auton et al. 2012) may be appropriate for chimps and bonobos, it is less appropriate for more distantly related taxa like Gorillas. In brief, using the map of (Kong et al. 2010), we computed the total genetic map length in cM and in bp, and scaled our maps to have the same expectation (i.e, same cM/MB), which yielded N_e_ estimates of 13428, 16781 and 19785 for bonobos, Nigerian chimpanzees and gorillas, respectively. The bonobo estimate falls within the population size estimated range of 11.9-23.8 in Table 1 of (Prado-Martinez et al. 2013). However, the gorilla and Nigerian chimpanzee size estimates are less than the estimated ranges of 26.8-53.5 for western lowland gorillas and 18.5-37 for Nigerian chimpanzees (Prado-Martinez et al. 2013). The low population size estimates are likely due to recent population size reductions in these species. These N_e_ values yielded adjusted rate estimates of ~1.193 cM/Mb for bonobo, Nigerian chimpanzee and gorillas.

Together with western chimp, the block boundaries for regions that are syntenic to human were intersected across the four non-human maps. This intersection resulted in a set of blocks that were designated as ‘multi-syntenic’ across all maps. The intersection was done using the bedTools program *multiIntersectBed* (Quinlan and Hall 2010). Next, blocks below 1Mb were removed from this set. Then, the map files were filtered to only include estimates within these blocks and blocks where all maps had a minimum of 90% representation. These reduced maps and multi-syntenic blocks were used for most comparative analysis presented in the results. This process was repeated using the multi-syntenic blocks to calculate mean rate estimates in intervals of 1 Mb, 500 kb, and 100 kb for each of the three maps generated here, the additional two human maps from HapMap, and the PanMap data. Figure 1 shows a boxplot of the mean values across all six maps using the 1 Mb binned dataset and the full 1 Mb binned dataset can be found in **Table S2**.

### Hotspot determination and sharing between populations

LDhot uses a composite-likelihood framework based on the work of (Hudson 2001; McVean et al. 2002). The Auton et al. (2012) implementation tests every 2 kb region (with a 1 kb increment) as a potential hotspot by analyzing the 200 kb region centered around the region of interest. Suppose the SNPs in the 200 Kb region are S = {s_1_,… s_n_}. A (composite) LRT statistic is calculated as:

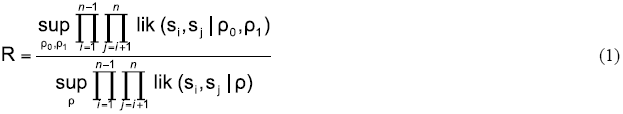

where lik (*s*_i_, *s*_j_ | ρ) is the 2-site likelihood described before (Hudson 2001; McVean et al. 2002), ρ_0_ is the background recombination rate and ρ_1_ is the recombination rate in the central 2 kb region.

Critical values for R are estimated from null simulations that assume a constant recombination rate across the region (i.e., ρ_0_ = ρ_1_). Auton et al. (2012) used ‘fixed S’ methodology for these simulations (Hudson 1993; Wall and Hudson 2001), with SNP locations fixed to be where SNPs appear in the actual data and ρ chosen to be equal to its estimated value (from LDhat). They tested each possible 2 kb region and identified those where the estimated p-value for R was < 0.01. Then, overlapping regions were merged to form a list of candidate hotspot regions. These regions were filtered to reduce the false positive rate by eliminating ones > 5 kb in size or with peak ρ estimate < 5/kb (estimated using LDhat). Auton *et al*. (2014) used a smaller window size (100 kb) but the same basic approach for identifying candidate regions. Instead of a size or peak ρ estimate filter though, they required each hotspot region to contain at least one 2 kb window where the estimated p-value for R was < 0.001. Our new approach here is to use a 20 kb window size, generate the same list of candidate regions, partition each region into non-overlapping 1 kb windows, and keep only those windows for which the average ρ estimate (using LDhat) is at least 5 times the genome-wide average rate.

### Comparisons between existing and newly identified hotspots

In order to compare recombination rates at hotspots identified here and in previous studies, HapMap hotspots were downloaded from ftp download site:

ftp://ftp.ncbi.nlm.nih.gov/hapmap/recombination/2006-10_rel21_phaseI+II/hotspots/. The HapMap hotspots represent a composite of hotspots identified in each population of HapMap and present in at least 2 or more populations. For PanMap, hotspots were downloaded from ftp site:

ftp://birch.well.ox.ac.uk/panMap/haplotypes/genetic_map/hotspots/. All hotspots were converted from species-specific coordinates to hg18 coordinates using the UCSC tool liftOver with minMatch=0.1. For the most part, only a small number of regions were lost in this conversion to hg18 (see Figure 2 for adjusted numbers of hotspots). Next, the coordinates for each full recombination map were re-scaled ± 20 kb relative to the center of each set of hotspots Then, a loess smoothing was applied to the rate estimates using the re-scaled coordinates across all hotspots in each map (Figure 2). Using *multiIntersectBed*, the Bed coordinates of each set of hotspots across Nigerian Chimp, Western Chimp and Bonobo were intersected revealing 0.885% hotspot overlap between Western and Nigerian Chimps, 1.247% overlap between Nigerian Chimp and Bonobo, and 0.843% overlap between Western Chimp and Bonobo. There were only 4 hotspots that overlapped between all three populations. Finally, the same method used to identify hotspots for the three recombination maps generated in this study was applied to the phased haplotype data retrieved from ftp site: ftp://birch.well.ox.ac.uk/panMap/haplotypes/VCF/. This resulted in a set of 9,993 hotspots in Western Chimp (as compared to 5,038 from the original set of Western Chimp hotspots). **Figure S7** shows the plot of this new set of Western Chimp hotspots with rates from all six compared maps (similar to Figure 2B). Using the same intersection approach as described above, the new set of Western Chimp hotspots have 1.453% overlap with Western Chimp, and 1.279% overlap with Bonobo, with 10 hotspots that overlapped between all three populations.

### Examining the relationship between PRDM9 binding motifs and hotspots

Previous work has shown higher recombination rates in PRDM9 predicted binding sites in humans, but not in western chimpanzees. To further investigate the importance of PRDM9 in localizing to population-specific hotspots, we downloaded the protein sequences of PRDM9 from previous studies (Berg et al. 2011; Auton et al. 2012; Schwartz et al. 2014). We then predicted binding motifs for the zinc-fingers of the protein sequence using http://zf.princeton.edu/ and the polynomial SVM model as described in (Persikov et al. 2009). For chimps and bonobos, we calculated position weight matrices for the three submotif regions (see Fig. S17 in Auton et al. 2012) and a recently described internal motif AnTTnnAnTCnTCC (see Fig. S2 in Schwartz et al. 2014). We chose to include all four *Pan* submotifs in our analysis across the *Pan* group. For gorillas, we used the internal motif CCnAnnCCTC identified in (Schwartz et al. 2014) and an additional submotif CTCnTCnTCnTC shown in **Figure S8**. Additionally, we downloaded six publicly available gorilla PRDM9 alleles from:

http://przeworski.c2b2.columbia.edu/index.php/softwaredata/. Based on the ten total PRDM9 alleles of gorilla PRDM9, we tested a few other submotifs early in our analysis, but none seemed to associate with recombination rates at hotspots. Further, the human-specific submotifs that have shown to associate with recombination occur near the end of the full PRDM9 protein sequence, therefore we chose the additional submotif shown in Figure S8 in gorilla to determine if the final few zinc fingers are more relevant for recombination rate association. To validate our approach, we also used the 13-bp submotif CCnCCnTnnCCnC identified in humans (Myers et al. 2010) and predominantly used in Europeans, as well as the 17-bp PRDM9 allele more common in Africans CCnCnnTnnnCnTnnC, described in (Hinch et al. 2011a). This analysis included eight submotifs with different combinations searched across six recombination maps (including the newly defined set of hotspots for western chimpanzee).

We used an alternative approach to Auton *et al*. (2012) to examine the prevalence of PRDM9 in explaining hotspot distribution. We computationally identified matched coldspots across the genome, matching for GC content allowing for no more than 2% deviation from GC content of the hotspot, distance to the hotspot (within 100kb) and length of the region. GC content was calculated from the unmasked version of the genome. We further required that the average recombination rate of the ‘cold’ region be lower than the genome average for the specific population and the peak recombination rate be below the cutoff used for identifying hotspots. We then extracted the fasta sequence for both the hotspots and coldspots from the masked version of the species-specific reference genome. Next, we used the software ‘fimo’ to identify potential binding sites within the fasta sequence of the hotspots and coldspots (Grant et al. 2011). In Table 1, we report the results for individual submotifs for each map and set of hotspots used here and the analysis summed across all submotifs. In addition to total number of predicted binding regions for both hotspots and coldspots, we summed up the number of non-zero regions for each to yield a proportion of regions with *any* suspected PRDM9 binding.

In order to explain why our results were different from those of Auton *et al*. (2012), we later performed a genome scan for each submotif. For this analysis, we split the genomes into 1kb regions and computed for each region recombination rate and GC content. We then ran fimo to get the predicted binding sites across each region. From these fimo results, we selected the top 5000 predicted binding sites across the genome to look for enriched recombination rates as compared to a null motif. We generated a null motif for each submotif by randomly shuffling the individual PWMs. This method is slightly different than previous studies which have simply replaced a single base in the PWM with another to generate a null motif. In **Figure S11**, we plot the results of our genome-wide survey of recombination rate enrichment at PRDM9 submotif predicted binding regions as compared to a null motif.

### Genomic distribution of recombination rates

To get cumulative distributions of recombination rates across each recombination map, the absolute physical and genetic distance for each interval was calculated, sorted relative to genetic distance, and summed to 1 for both physical and genetic distance values across the full dataset (*5*). For this analysis, we chose to plot the cumulative distribution using the R package *ineq*, where it is referred to as the Lorenz curve (**Figure 3A**). The diagonal shows the expectation under a uniform distribution of recombination rates, and the deviation from the diagonal represents increasing bias in the distribution towards recombination hotspots. From these data, we calculated the Gini coefficient for all six maps (**Figure 3B**).

### Broad-scale comparisons

Pairwise nucleotide divergence between each population was taken from Suppl. Table 5.2 of Prado-Martinez *et al* 2013. To compare each map, the 1 Mb rate estimates from the multi-syntenic regions as defined between all six maps were fitted to a regression in R and the Pearson correlation coefficient between each pairwise comparison was computed.

Using the 1 Mb binned multi-synteny dataset, we examined variation in the skew of recombination typically observed at chromosome ends (Serre et al. 2005). We split the data into p and q arms of each chromosome and removed arms less than 30 Mb to make sure that a minimum of 15 Mb for each arm could be independently attributed to centromeric and telomeric skews. We then adjusted the distance to be relative to the chromosome end and binned the data into 1 Mb bins. To normalize the rate estimates on each arm, we calculated the percent difference of each 1 Mb bin from the mean rate for the whole arm. We then plotted the skew at telomeres (**Figure S15A**) and centromeres (**Figure S15B**) in 1 Mb bins for the first and last 25 Mb relative to the chromosome ends for all six comparative maps.

In addition to skews in recombination due to chromosomal location, we examined how largescale changes in chromosomal structure impacted recombination rates in great apes. We examined the chromosome 2 junction across great apes and the translocation of human chromosomes 5 and 17 in gorillas.

### Multiple Linear Regression Analysis

From the UCSC genome browser, we downloaded: three classes of repeats, Repeating Elements (v. 3.2.7), Simple Tandem Repeats, Microsatellites, two classes of functional elements, exons from the CCDS project and phastCons elements from the 28-way placental mammals alignments, and three Encode annotations pertaining to gene activity - Transcription-associated, Active Promotor, and Strong Enhancer - from the Genome Segments track (from GM12878, combined Segway + ChromHMM). The Encode annotations were converted from hg19 to hg18 coordinates using the UCSC liftOver tool set to the default parameters.

We calculated nucleotide diversity (π), divergence, ancestral GC content and GC-flux using the ancestral-sequence inferred for the common ancestor of humans and orangutans using the same methodology as described in Prado-Martinez et al. (2013). Briefly, to ensure that orthologous bases were used, we used the read-mappings of all of the great ape resequencing data to the hg18 reference genome, and we used a merged callablility mask which ensures that the same nucleotides, which are callable in all taxa, are used in our analyses. Using these data, we computed pairwise diversity (π), the mean divergence rate to the ancestor of humans and orangutans (inferred in Prado-Martinez et al. 2013), and GC flux on substitutions, as well as the ancestral GC-content. While the use of the hg18-reference genome, as opposed to species-specific genomes, may induce reference biases in our calculations, these biases are inherent with the inclusion of bonobos in our analysis, which was mapped to the chimpanzee reference genome. Further, the use of the same reference genome across taxa ensures that the exact same nucleotides are compared in our analyses.

Annotations were computed on the base-wise unions of each of the annotations as appropriate using BedTools (Quinlan and Hall 2010). Specifically, each locus was annotated with the fraction of bases that compose the above categories. Local recombination rates were computed using the species/population-specific genetic maps at varying window sizes (100kb, 500kb and 1Mb nonoverlapping loci) taken from the multisyntenic regions described earlier.

Performing an ordinary least-squares regression on whole genome data violates several model assumptions, including autocorrelation (e.g., linkage-disequilibrium), heteroscadasticity, and multicollinearity. Modern regression techniques can address some of these issues through the use of robust regression. Using the rlm command in R, we performed a robust linear regression to model the local recombination rate as a linear function of the annotations described above. Robust regression does an iterated reweighted linear least squares to down-weight the impact that outliers have on the coefficients. As our independent variables are intrinsically correlated with each other (e.g., GC content and exonic content), we computed variance inflation factors (VIFs) to assess how problematic these associations are for our analyses. Across our linear models, the maximum VIF was less than 5 (a VIF of 5 indicates that our standard errors are inflated by a factor of 5), indicating that while multicollinearity is present, our inferences should be relatively robust nonetheless. Examination of the coefficient + residual plots showed relationships in the variables that were largely linear in nature, suggesting that a linear model may be appropriate for this analysis. As heteroscedasticity and autocorrelation artificially reduce our variance estimates, we used the sandwich package in R to get heteroscedasticity-and autocorrelation-consistent variance/covariance matrix estimates using the vcovHAC and coeftest functions. Beta (standardized) coefficients were computed by converting of the variables to have mean 0 and variance 1 using the scale function in R, prior to running the linear regression, which allows magnitude of the coefficients to be compared within and between models. Model coefficients were visualized with the coefplot function in R.

As many categories of functional elements are included in our model, including phylogenetically conserved elements, gene definitions, as well as annotations from the Encode project, the correlation between local recombination rate and divergence points to the possibility that these elements are unannotated and have large (combined) effects, consistent with recent predictions (McManus et al. *In press*). Further, linkage to selective elements in the ancestor of humans and orangutans can also induce a correlation between recombination and divergence (Begun and Aquadro 1992); however, all taxa are compared against the same ancestor.

## Data Access

We plan to deposit all data associated with this manuscript onto Dryad for dissemination. We also plan to make all scripts generated to for analysis completed here available on Github.

## Acknowledgments

This work was supported by the National Human Genome Research Institute of the National Institutes of Health under award number R01_HG005226 to MFH and JDW and National Institute Of General Medical Sciences of the National Institutes of Health under Award Number F32GM101744 to LSS. The content is solely the responsibility of the authors and does not necessarily represent the official views of the National Institutes of Health. AEW is supported by National Science Foundation Graduate Research Fellowship Grant DGE-1143953. Computations for this study were performed on the QB3 cluster at UCSF.

## Author contributions

LSS, AEW and JDW performed analyses. LSS and JDW wrote the manuscript. JMK, JLK and CDB analyzed raw data, performed variant calling, and initial filtering. JMK, JLK, KRV, MFH and CDB performed relatedness analysis on GAGP samples for inclusion in this study. CDB, KM, and MFH gave ongoing feedback and conceptual advice. All authors contributed to interpretation of the results and provided critical review and intellectual content on the draft manuscript.

